# Mapping the unique and shared functions of oncogenic KRAS and RIT1 with proteome and transcriptome profiling

**DOI:** 10.1101/2020.04.10.030460

**Authors:** April Lo, Kristin Holmes, Filip Mundt, Sitapriya Moorthi, Iris Fung, Shaunt Fereshetian, Jackie Watson, Steven A. Carr, Philipp Mertins, Alice H. Berger

## Abstract

Aberrant activation of RAS oncogenes is prevalent in lung adenocarcinoma, with somatic mutation of *KRAS* occurring in ∼30% of tumors. Recently, we identified somatic mutation of the RAS-family GTPase *RIT1* in lung adenocarcinoma, but relatively little is known about the biological pathways regulated by RIT1 and how these relate to the oncogenic KRAS network. Here we present quantitative proteomic and transcriptomic profiles from *KRAS*-mutant and *RIT1*-mutant isogenic lung epithelial cells and globally characterize the signaling networks regulated by each oncogene. We find that both mutant KRAS and mutant RIT1 promote S6 kinase, AKT, and RAF/MEK signaling, and promote epithelial-to-mesenchymal transition and immune evasion via HLA protein loss. However, KRAS and RIT1 diverge in regulation of phosphorylation sites on EGFR, USO1, and AHNAK proteins. The majority of the proteome changes are related to altered transcriptional regulation, but a small subset of proteins are differentially regulated by both oncoproteins at the post-transcriptional level, including intermediate filament proteins, metallothioneins, and MHC Class I proteins. These data provide the first global, unbiased characterization of oncogenic RIT1 network and identify the shared and divergent functions of oncogenic RIT1 and KRAS GTPases in lung cancer.

## Introduction

Somatic mutation of the *KRAS* proto-oncogene is a prevalent feature of human cancers, particularly in lung adenocarcinomas where *KRAS* is mutated in up to 30% of tumors. Cancer-associated variants such as G12V and Q61H alter the normal regulation of KRAS GTPase activity by disrupting GTP hydrolysis activity or physical interaction with GTPase-activating proteins (GAPs)^1,2^, resulting in heightened downstream cellular signaling through the canonical RAS effector pathways RAF/MEK and PI3K/AKT as well as others. Following the discovery of cancer-associated RAS mutations in the 1980s^3,4^, thousands of studies have delineated the critical pathways involved in RAS-mediated cellular transformation, metastasis, and metabolism.

Recently, another RAS-family GTPase gene, *RIT1*, was found to harbor somatic mutations in lung cancer^5^ and myeloid leukemias^6^. Interestingly, germline *RIT1* mutations are found in families with Noonan Syndrome, a developmental “RAS”-opathy involving altered craniofacial morphology and cardiac abnormalities^7^, and which can also be caused by germline mutations in *KRAS* itself or other RAS-pathway genes such as *SOS1, SOS2, LZTR1* and *SHOC2* (https://omim.org/). In cancer and development, *RIT1* mutations are found in cases that lack canonical *KRAS* mutations, suggesting that *RIT1* may impart the same phenotypes conferred by activation of RAS.

Prior studies have characterized the role of RIT1 in neural development^8^ and we and others have described the role of mutant RIT1 in cellular transformation^5,9,10^, knowledge of the function of cancer- and Noonan-associated *RIT1* variants is relatively limited. Unlike *KRAS, RIT1* mutations are rarely observed near the critical glycine residues involved in GTP hydrolysis (e.g. G12 and G13 in KRAS or G30 and G31 in RIT1). Instead, *RIT1* mutations occur most frequently near the switch II domain, also targeted by Q61 KRAS variants (**Fig. 1a**). Nevertheless, these mutations may enhance GTP-bound levels of RIT1^11,12^. The molecular consequences of RIT1 switch II domain mutations may additionally be linked to the loss of RIT1’s physical interaction with LZTR1, a ubiquitin-conjugating enzyme responsible for degradation of RIT1^11^. Cancer- and Noonan-associated RIT1 variants lose the ability to interact with LZTR1 and consequently are highly overexpressed, resulting in increased signaling activity through the RAF/MEK pathway^11^.

**Figure 1.**
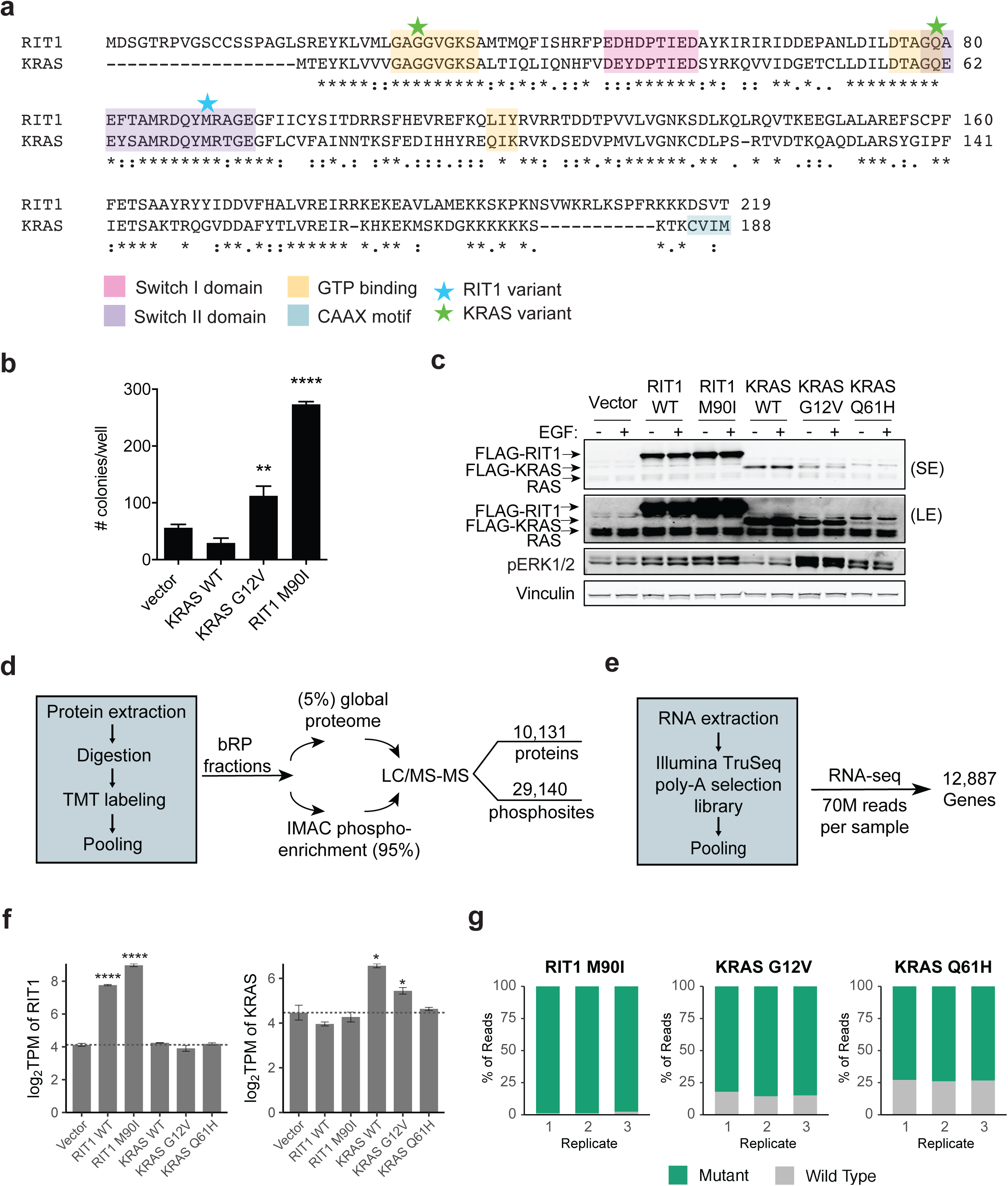
Comparative multi-omic profiling of KRAS- and RIT1-mutant human lung epithelial cells. **a**, Protein alignment of KRAS-4B (Uniprot #P01116-2) and RIT1 Isoform 1 (Uniprot #Q92963-1) generated by ClustalW2^61^. Stars indicate the position of the RIT1^M90I^ or KRAS^G12V^ and KRAS^Q61^ variants used in this study. Asterisks indicate fully conserved residues. Colons indicate strongly conserved residues. Periods indicate weakly conserved residues. **b**, Soft agar colony formation assay of isogenic AALE human lung epithelial cells. **, p < 0.01; ****, p<0.0001 by two-tailed t-test. **c**, Western blot using anti-RAS and anti-RIT1 antibodies (top panels), or antibodies against phosphorylated ERK1/2 or vinculin (loading control). SE = short exposure, LE = long exposure. Isogenic AALE cells were cultured in the presence or absence of EGF for 12 hours. **d**, LC-MS/MS workflow for generation of proteome and phosphoproteome profiles. bRP, basic reverse phase chromatography. IMAC, immobilized metal affinity chromatography. **e**, Workflow for Illumina RNA-seq analysis. **f**, mRNA quantification in transcripts per million (TPM) showing mean ± SD of RIT1 (left) or KRAS (right) in isogenic AALE cells, n = 3 per cell line. *, p < 0.05; ****, p < 0.0001 by two-tailed Student’s t-test compared to vector control cells. **g**, RNA-seq quantification of variant allele expression. Data shown is the percentage of reads at the M90I, G12V, or Q61H variant site for the variant allele or wild-type allele.

Prior studies of RIT1 function focus on candidate cellular signaling pathways based on RIT1’s homology to KRAS. To our knowledge, unbiased mapping of downstream RIT1-regulated pathways has not been performed to date. Here we sought to broadly describe the proteome, phosphoproteome, and transcriptome changes induced by wild-type RIT1 and RIT1^M90I^, a cancer- and Noonan-associated variant, and to compare these changes to those induced by oncogenic KRAS variants. With a particular interest in the consequences of RIT1^M90I^ in lung cancer, we profiled the effects of RIT1^M90I^ mutation in AALE cells, a non-transformed, immortalized, human lung epithelial cell line^13^.

By comparing the downstream pathways regulated by oncogenic KRAS and RIT1, we uncover previously unknown consequences of RIT1 activation, such as induction of the epithelial-to-mesenchymal transition (EMT) and post-translational regulation of HLA protein expression. In addition, we uncover additional functional differences between KRAS and RIT1 including a distinct and unique role of KRAS mutants in regulation of EGFR and USO1 phosphorylation. These data provide the first systems-level view of RIT1 and RIT1^M90I^ function.

## Results

### Multi-omic profiling of RIT1- and RAS-transformed human lung epithelial cells

We previously demonstrated that RIT1^M90I^ and other cancer-associated RIT1 variants can promote cellular transformation of NIH3T3 cells in vitro and in vivo^5^. To determine whether RIT1^M90I^ was capable of transforming human lung epithelial cells, we expressed mutant RIT1 or KRAS in the human lung epithelial cell line, AALE. Similar to our previous findings in rodent cells, both RIT1^M90I^ and KRAS^G12V^ enabled AALE cells to form colonies in soft agar (**Fig. 1b**).

The canonical function of oncogenic RAS variants is the downstream activation of the RAF/MEK/ERK cellular signaling cascade^14^, and RIT1 shares the ability to bind C-RAF and induce transcription of ERK target genes activity^11^. To determine if such regulation is active in AALE cells, we stably expressed wild-type RIT1 or KRAS, or the mutant forms RIT1^M90I^, KRAS^G12V^, and KRAS^Q61H^ in AALE cells. KRAS^Q61H^ was included since this mutant more closely resembles the switch II domain mutants observed in *RIT1* in cancer (**Fig. 1a**). RIT1^M90I^, KRAS^G12V^, and KRAS^Q61H^ all enhanced ERK phosphorylation compared to their respective wild-type protein or vector control (**Fig. 1c**). Interestingly, wild-type RIT1 overexpression also modestly enhanced ERK phosphorylation whereas wild-type KRAS suppressed basal ERK phosphorylation.

To systematically characterize the signaling networks perturbed by mutant RIT1 and KRAS in lung cancer, we expressed each variant in AALE cells and performed both RNA-seq and deep proteome and phosphoproteome profiling by liquid chromatography tandem mass spectrometry (LC-MS/MS). Following trypsin digestion, peptides were labeled with tandem mass tag (TMT) reagents in two overlapping 10-plex sets for relative quantification of proteome and phosphopeptides by LC-MS/MS (**Fig. 1d and Supplementary Fig. 1a-b**). Following basic reverse phase chromatography, fractions were either directly subjected to LC-MS/MS for total proteome quantification, or subjected to immobilized metal affinity chromatography (IMAC) to enrich for phosphopeptides and then subjected to LC-MS/MS, or. In total, we identified 10,131 proteins, 9002 of which were detected and quantified in every sample, and 29,140 phosphopeptides, 12,325 of which were identified in common in every sample (**Supplementary Tables 1 and 2 and Supplementary Files 1 and 2**).

In parallel, we generated deep transcriptome profiles of the same isogenic cell lines. Transcriptome profiling was performed in triplicate on the Illumina NovaSeq platform to a median read-depth per replicate of 70.1 million reads (**Fig. 1e, Supplementary Table 3 and Supplementary Fig. 1e**). No compensatory feedback regulation of RIT1 to KRAS or vice versa was observed (**Fig. 1f**). Despite relatively low protein expression of KRAS variants in the AALE lines (**Fig. 1c**), the majority of KRAS transcripts in each isogenic cell line corresponded to G12V or Q61H variants, respectively, with 84.1% of reads harboring the G12V variant in KRAS^G12V^ cells, and 73.3% of reads corresponding to the Q61H allele in KRAS^Q61H^ cells (**Fig. 1g**). As expected, known KRAS-regulated gene sets were strongly up- and down-regulated in KRAS-mutant cells (**Supplementary Fig. 1d**).

### Multi-omic profiling identifies global similarity between signaling regulated by RIT1^M90I^ and oncogenic KRAS

Differentially abundant proteins were identified by comparison to the vector control cells using a two-tailed moderated t-test (**Fig. 2a**). Selected proteins observed to be significantly modulated by LC-MS/MS were cross-validated by Western blot. FOSL1, also known as FRA1, is a basic leucine zipper transcription factor in the FOS family^15^. Activation of RAS is known to promote transcriptional upregulation and protein stabilization of FOSL1^16,17^. By LC-MS/MS, FOSL1 was markedly overexpressed in KRAS^G12V^, KRAS^Q61H^, and RIT1^M90I^-mutant cells compared to wild-type cells or vector control cells (**Fig. 2b**). Consistently, Western blot of independently-derived AALE isogenic lines demonstrated greater abundance of FOSL1 in KRAS- or RIT1-mutant cells compared to wild-type expressing cells (**Fig. 2b**). TXNIP is an inhibitor of thioredoxin involved in both redox regulation and glucose metabolism^18,19^. Prior literature identified HRAS^G12V^-induced suppression of TXNIP transcription and protein translation^20,21^. TXNIP was among the top down-regulated proteins in KRAS- and RIT1-mutant proteomes, and was decreased in Western blot analysis of independently derived cells (**Fig. 2c**). These validation data demonstrate the utility of LC-MS/MS to describe protein expression changes and additionally suggest the mechanism of RAS-mediated modulation of FOSL1 and TXNIP is shared with RIT1^M90I^.

**Figure 2.**
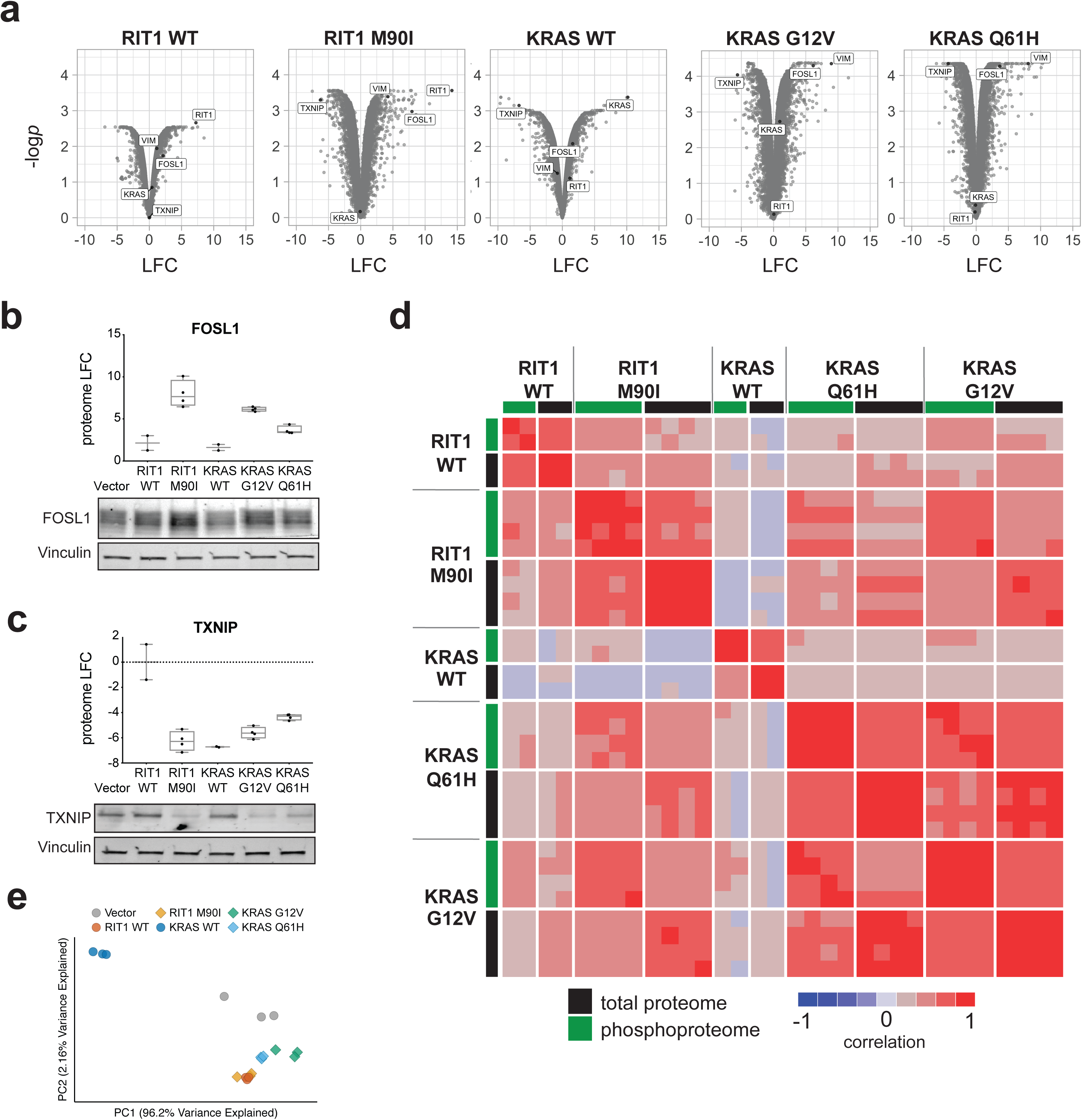
Quantitative proteome and transcriptome profiling identifies similarity in RIT1^M90I^-mutant and KRAS-mutant signaling networks. **a**, Volcano plots of global proteome data from isogenic AALE cells showing the log_2_(fold change) (“LFC”) in protein abundance in each cell line compared to vector control cells. The y-axis displays the negative log_10_(*p* value) calculated from a one sample moderated t-test with multiple hypothesis correction by the Benjamini-Hochberg method. **b**, Western blot validation of FOSL1 increased protein abundance in RIT1- and KRAS-mutant cells. The chart shows the LFC of FOSL1 as determined by LC-MS/MS. Western blot below shows FOSL1 abundance or Vinculin (loading control). **c**, Western blot validation of TXNIP protein abundance in RIT1- and KRAS-mutant cells. The chart shows the LFC of TXNIP as determined by LC-MS/MS. **d**, Correlation heatmap showing pairwise Pearson and Spearman correlations of each proteome and phosphoproteome replicate to every other replicate. To enable correlation of proteome with phosphoproteome, phosphosites were collapsed to the protein level by taking the median of all phosphosites for each protein. **e**, Principal component analysis (PCA) of RNA-seq data. Circles correspond to control vector or wild-type replicates. Diamonds correspond to RIT1- or KRAS-mutant profiles.

Next we compared the global effects of RIT1^WT^ and RIT1^M90I^ to that of KRAS^WT^ and KRAS variants. Proteome and phosphoproteome data from RIT1^M90I^-expressing cells were highly correlated with KRAS^G12V^ and KRAS^Q61H^ profiles, suggesting largely similar downstream consequences (*r* = 0.70-0.80 and 0.72-0.75 for proteome and phosphoproteome, respectively; **Fig. 2d**). Despite differences in KRAS protein abundance, KRAS^G12V^ and KRAS^Q61H^ proteomes and phosphoproteomes were highly correlated (proteome *r* = 0.85 and phospho *r* = 0.79; **Fig. 2d**). In contrast, wild-type KRAS replicates were the most divergent of all profiles, showing limited correlation to either the KRAS-mutant profiles or RIT1 profiles.

A recent study found that RIT1 variants, including M90I, may function by relieving negative regulation of RIT1 by a LZTR1-dependent proteasomal degradation mechanism^11^. Accordingly, overexpression of wild-type RIT1 should largely phenocopy expression of RIT1^M90I^. Consistent with this idea, RIT1^WT^ cells more closely resembled both RIT1^M90I^ and KRAS-mutant cells than KRAS^WT^ cells (**Fig. 2d**). These data highlight a critical divergence between KRAS and RIT1: expression of wild-type KRAS is not capable of activating downstream oncogenic pathways, whereas expression of wild-type RIT1 in part resembles activation of RIT1 or KRAS by mutation. We confirmed this observation in a principal component analysis of transcriptome data, which further revealed a high degree of similarity between RIT1^WT^ and RIT1^M90I^-regulated gene expression (**Fig. 2e and Supplementary Table 4**).

### Oncogenic RIT1 promotes epithelial-to-mesenchymal transition

To identify the downstream pathways regulated by oncogenic KRAS and RIT1, we performed gene set overlap analysis using MSigDB Hallmark Pathway gene sets^22^ (**Fig. 3a**). The epithelial-to-mesenchymal transition (EMT) pathway was the most significant gene set enriched among up-regulated proteins for both KRAS^G12V^/KRAS^Q61H^ and RIT1^WT^/RIT1^M90I^ cell lines. EMT is a cellular transdifferentiation process promoted by cell-extrinsic signaling proteins and orchestrated by activation of transcription factors such as Twist, Snail, and Zeb family transcription factors^23^. It has long been observed that oncogenic RAS proteins, including KRAS mutants, promote EMT. An EMT-signature is associated with KRAS dependence^24^, which has been functionally linked to activation of FOSL1^25^. Interestingly, we find both RIT1^M90I^ and KRAS^G12V^/KRAS^Q61H^ are capable of promoting expression changes of key EMT markers, including up-regulation of Vimentin, N-Cadherin, and Fibronectin, and downregulation of Keratin 19 (**Fig. 3b and Supplementary Fig. 2a**). Although canonical EMT transcription factors Snail (SNA1) and Slug (SNA2) were not detected by proteomic analysis, transcriptomes from RIT1- and KRAS-mutant cells showed increased activity of these EMT transcription factors as determined by ChEA3 transcription factor enrichment analysis (**Fig. 3c-d and Supplementary Fig. 2b**). To our knowledge, this is the first demonstration of mutant RIT1 promoting EMT in any cell type.

**Figure 3.**
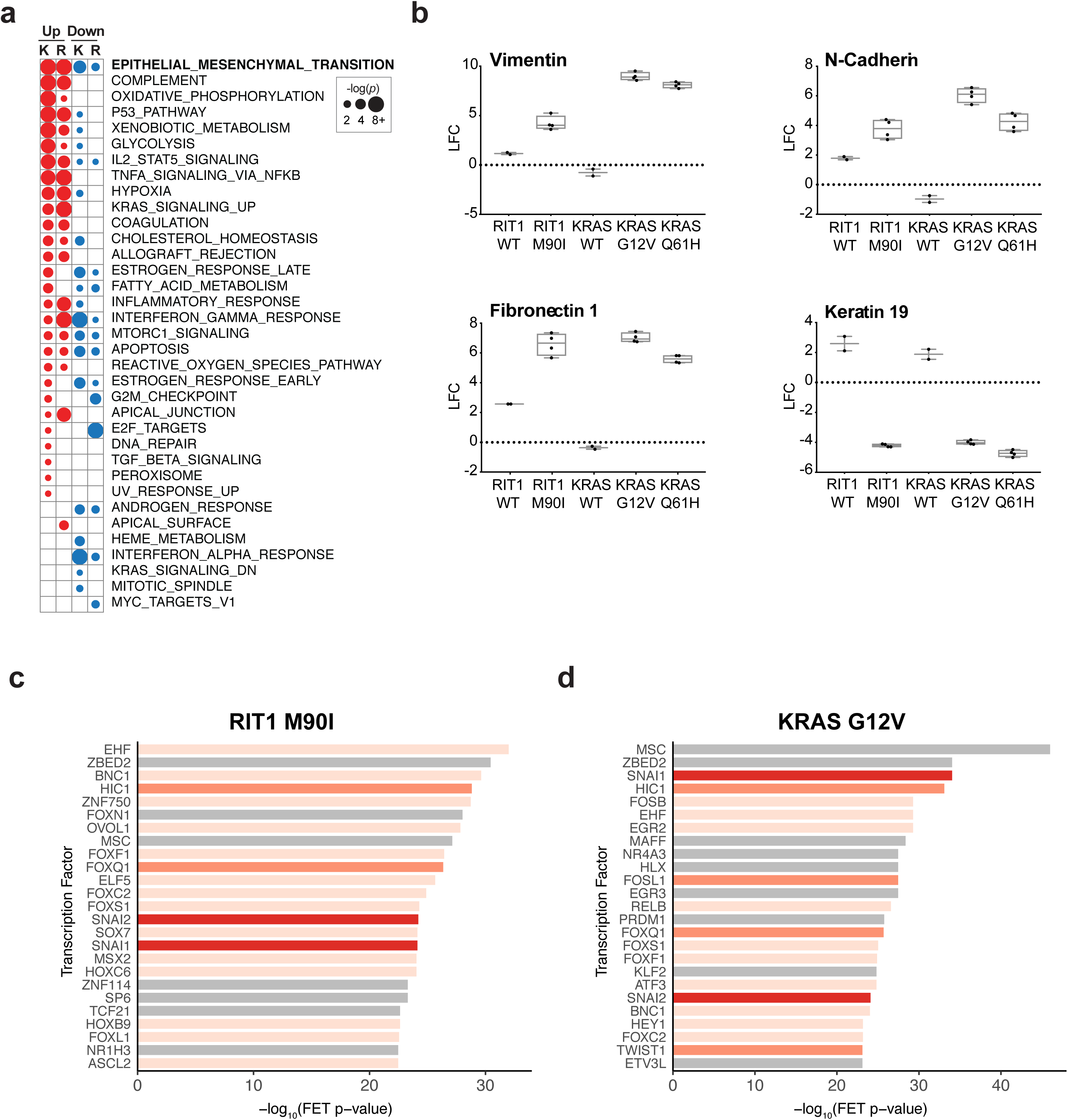
RIT1^M90I^ promotes epithelial-to-mesenchymal (EMT) transition. **a**, Gene set overlap analysis of up-regulated (“Up”; LFC>2) and down-regulated (“down”; LFC<-2) proteins using MSigDB Hallmark Pathways^22^. “K” and “R” indicate analysis based on mean LFC of KRAS^G12V^/KRAS^Q61H^ cells or RIT1^WT^/RIT1^M90I^ cells, respectively. Circle size corresponds to the *p* value of gene set overlap analysis determined by MSigDB. **b**, LFC of protein abundance of EMT marker genes as determined by LC-MS/MS, relative to vector control cells. **c**, Transcription factor target enrichment analysis of differentially expressed genes in RIT1^M90I^-mutant cells using Enrichr libraries. FET, Fisher’s exact test. Red = Snail family. Orange = confirmed EMT genes in dbEMT^59^. Pale orange = EMT-associated genes in literature. **d**, Enrichr analysis of KRAS^G12V^- mutant proteome data. Annotation is the same as in **c**.

### Oncogenic KRAS and RIT1 suppress Class I MHC expression via a post-transcriptional mechanism

Among the top suppressed proteins with differential abundance in both mutant KRAS and RIT1^M90I^ cells, were major histocompatibility complex (MHC) proteins. Class I MHC proteins HLA-A, HLA-B, HLA-C, and HLA-F were potently suppressed by KRAS^G12V^, KRAS^Q61H^, and RIT1^M90I^ (**Fig. 4a-b and Supplementary Fig. 3a**). Recently there has been a renewed interest in expression of immune modulatory proteins in cancer due to the appreciation of the potent role of the immune system in shaping cancer evolution. Further understanding the regulation of HLA expression in cancer is particularly critical in metastatic *KRAS*-mutant lung adenocarcinoma, where chemotherapy combined with immune checkpoint blockade is often used in the first-line setting.

**Figure 4.**
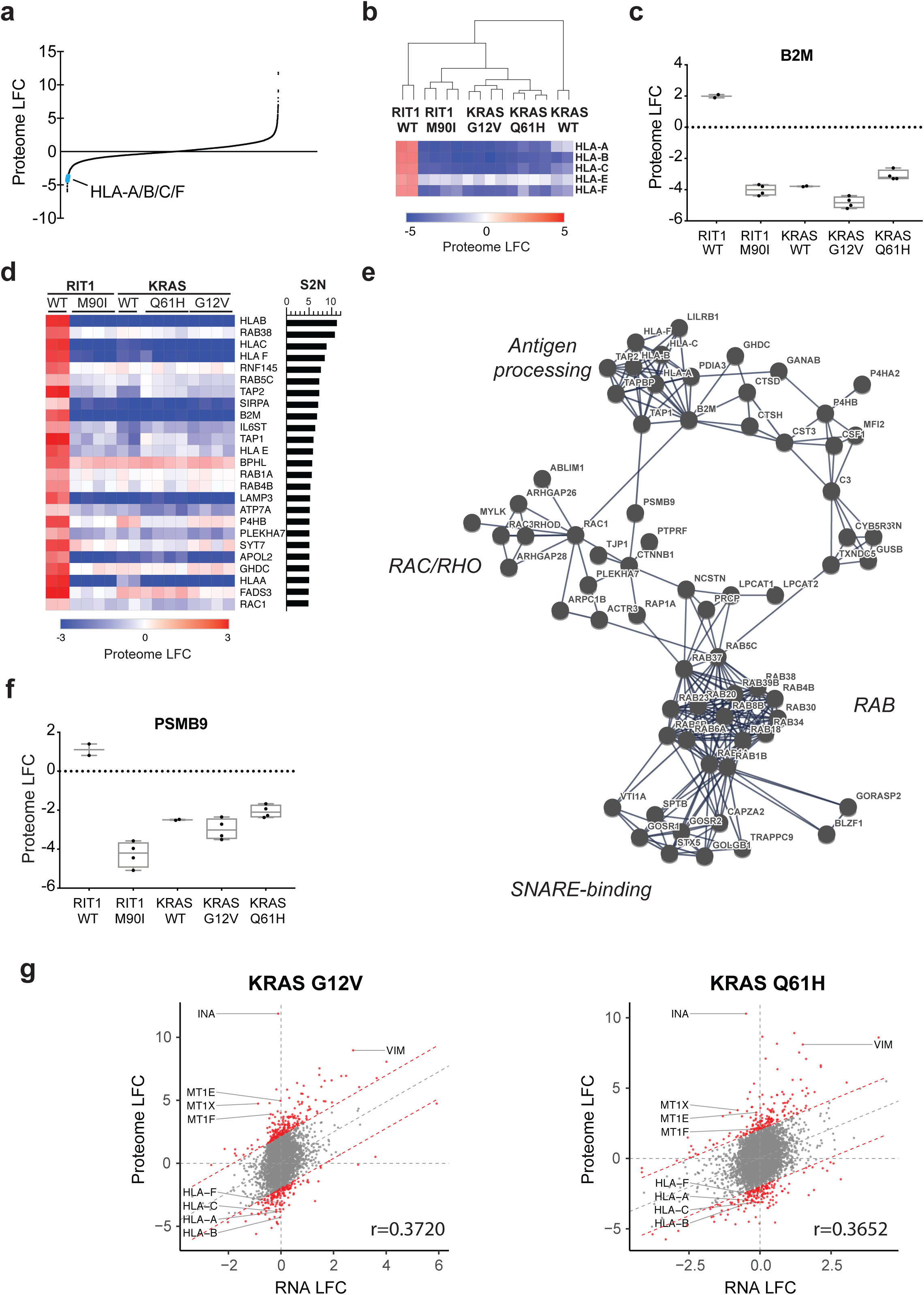
RIT1- and KRAS-mutant cells suppress Class I MHC expression via global loss of antigen processing and presenting machinery. **a**, Rank plot of all protein abundance changes in KRAS^G12V^-mutant cells compared to vector control, generated by LC-MS/MS. HLA-A,-B,-C, and -F proteins are labeled in blue. **b**, Heat map showing HLA protein abundance in each global proteome replicate. Replicates were clustered by unsupervised hierarchical clustering using all detected proteins. **c**, Protein abundance of B2M in LC-MS/MS data. **d**, Top differentially abundant proteins between RIT1^WT^ cells and all other cell lines. Proteins are ranked by the signal-to-noise (S2N) statistic, shown in the bar chart at the right. **e**, StringDB^62^ network analysis of proteins with S2N>2.5 in analysis shown in **d**. The network was significantly more connected than expected by chance (*p* < 1e-16). Disconnected nodes, single connected nodes, and disconnected clusters have been removed from the visualization. Edges represent high confidence interaction scores (>0.9) and network edge thickness indicates the strength of data support from all StringDB active interaction sources. **f**, Protein abundance of PSMB9 in LC-MS/MS data. **g**, Global proteome-transcriptome correlation analysis. A dashed diagonal line displays the linear regression generated by comparing the LFC of each gene in the transcriptome to its respective protein LFC in the proteome. The resulting Pearson correlation coefficient (*r*) is shown. Genes outside the 95% prediction interval are plotted in red, and include HLA genes, metallothioneins, and intermediate filament proteins Vimentin (VIM) and alpha internexin (INA).

Class I MHC genes *HLA-A, HLA-B*, and *HLA-C* harbor loss-of-function mutations in cancer^26^, demonstrating selective pressure to lose MHC function during tumorigenesis. Both MHC expression loss and upregulation of the immune suppressive protein PD-L1 enable tumor evasion of T-cell recognition of aberrant cancer cell proteins^27^. Moreover, expression loss of HLA proteins or B2M, another MHC Class I complex protein, is associated with resistance to immunotherapy in cancer^28^. We found that RIT1^M90I^, KRAS^G12V^, and KRAS^Q61H^ cells all promoted loss of B2M protein abundance in addition to HLA protein loss (**Fig. 4c**).

Class I MHC expression is known to be dynamically regulated by upstream signals controlled by interferon gamma exposure, NF-kB signaling, and chromatin regulators such as EZH2^29,30^. Each of these mechanisms involves transcriptional regulation of class I MHC genes. However, there were no transcriptional differences in HLA genes in the KRAS-mutant and RIT1-mutant cells nor were any transcriptional differences observed in the upstream regulators of MHC Class I expression NLRC5 and IRF1 and IRF2 (**Supplementary Fig. 3b**). Moreover, we excluded the possibility that lentiviral transduction or expression of a foreign antigen was responsible for the HLA suppression, because HLA protein expression was maintained or enhanced in RIT1^WT^-expressing cells as well as vector control cells, which express the Renilla luciferase gene.

To identify the possible mechanism of RIT1^M90I^- and KRAS-mediated MHC suppression, we identified other proteins that, like HLA, were upregulated in RIT1^WT^ cells but suppressed in RIT1-mutant and KRAS-mutant cells (**Fig. 4d**). This analysis revealed the pervasive downregulation of the Rab-mediated ER/Golgi vesicle-trafficking pathway that controls MHC Class I processing and presentation as well as the MHC Class I complex proteins themselves (**Fig. 4e**). In addition, expression of the proteasomal subunit PSMB9 correlated with loss of the MHC processing machinery (**Fig. 4f and Supplementary Fig. 3c**). Loss of PSMB9, also known as LMP2, has been previously linked to loss of MHC expression after oncogenic transformation^31^. We conclude that RIT1^M90I^ and KRAS^G12V^/KRAS^Q61H^ suppress MHC Class I expression through a post-transcriptional mechanism possibly involving PSMB9. Further investigation of MHC Class I expression loss driven by these oncogenic RIT1 and KRAS is critical to better understand the role of RAS and RIT1 signaling on immune evasion in cancer.

The identification of a major class of proteins regulated at the post-transcriptional level in RIT1- and KRAS-transformed lung epithelial cells brought to our attention the possibility of other post-transcriptional regulation by RIT1 and KRAS. Indeed, oncogenic RAS signaling profoundly alters cap-dependent translation via activation of the p90 ribosomal S6 kinases (RSKs)^32^ and PI3K/mTOR^33^, so differential protein translation could significantly contribute to altered protein abundance in RAS-transformed cells. To determine whether there were other protein classes in addition to MHC Class I proteins with significant post-transcriptional regulation, we performed a global correlation analysis of the transcriptome and proteome. Significant linear correlations between transcript and protein abundance were observed for RIT1 and KRAS variants, with the correlation highest for cells expressing mutant KRAS^G12V^ (*r* = 0.3725) or KRAS^Q61H^ (*r* = 0.3620) (**Fig. 4g**). While expression of the majority of genes were correlated at the RNA and protein levels, the metallothionein protein family including MT1E, MT1F and MT1X was highly upregulated in the proteome but not transcriptome of KRAS-mutant cells (**Fig. 4g**). In addition, intermediate filament proteins were also substantially regulated post-transcriptionally; both alpha-internexin (INA) and vimentin (VIM) were expressed more highly in the proteome than expected from RNA-seq data (**Fig. 4g**). These data highlight the utility of LC-MS/MS to identify protein abundance changes that would not be predicted from transcriptome analysis.

### Phosphoproteome profiling illuminates shared and unique signaling by RIT1 and KRAS

Protein phosphorylation is a reversible and dynamic mechanism of intracellular signaling that enables rapid intracellular transduction of signals controlling cell proliferation, survival, and metabolism. Although both RIT1 and KRAS act as GTPase switches, they both stimulate activation of cellular protein kinases such as BRAF. We therefore evaluated protein phosphorylation regulated by wild-type and mutant RIT1 and KRAS. Phosphosite abundance was expressed as a relative abundance normalized to the total protein abundance for each phosphoprotein. Unsupervised hierarchical clustering of the phospho-signatures identified the RIT1^M90I^ phosphoproteome as most similar to KRAS^G12V^ and KRAS^Q61H^ phospho-signatures (**Supplementary Fig. 4a**). We performed Kinase-Substrate Enrichment Analysis (KSEA^34^), which uses kinase-substrate pairings from PhosphoSitePlus^35^ and NetworKIN^36^ to identify differential phosphorylation of kinase-substrate families (**Supplementary Table 5**). These data further confirmed the similarity in phosphorylation state between RIT1-mutant and KRAS-mutant cells. The top kinases with increased substrate phosphorylation in RIT1-mutant and KRAS-mutant cells were ribosomal S6 kinase (RPS6KA1), Protein kinase C (PRKCA), AKT1, and MAPKAPK2 (**Fig. 5a-c, Supplementary Fig. 4b-e**, and **Supplementary Table 5**). The levels of phosphorylation of RPS6KA1 and MAPKAPK2 substrates were enhanced most strongly in the mutant cells and less in RIT1 WT and KRAS WT-expressing cells (**Fig. 5b-c**). Substrates of Aurora kinase B and CDK1 and PAK1 were suppressed in RIT1- and KRAS-mutant cells (**Fig. 5a** and **Supplementary Fig. 4b-c and Supplementary Fig. 4f)**. Although the total phosphorylation of each substrate reflects the balance between kinases and phosphatases in the cell, these data suggest that RIT1^M90I^, like oncogenic KRAS, can activate the canonical RAS effector pathways involving S6 kinase and AKT.

**Figure 5.**
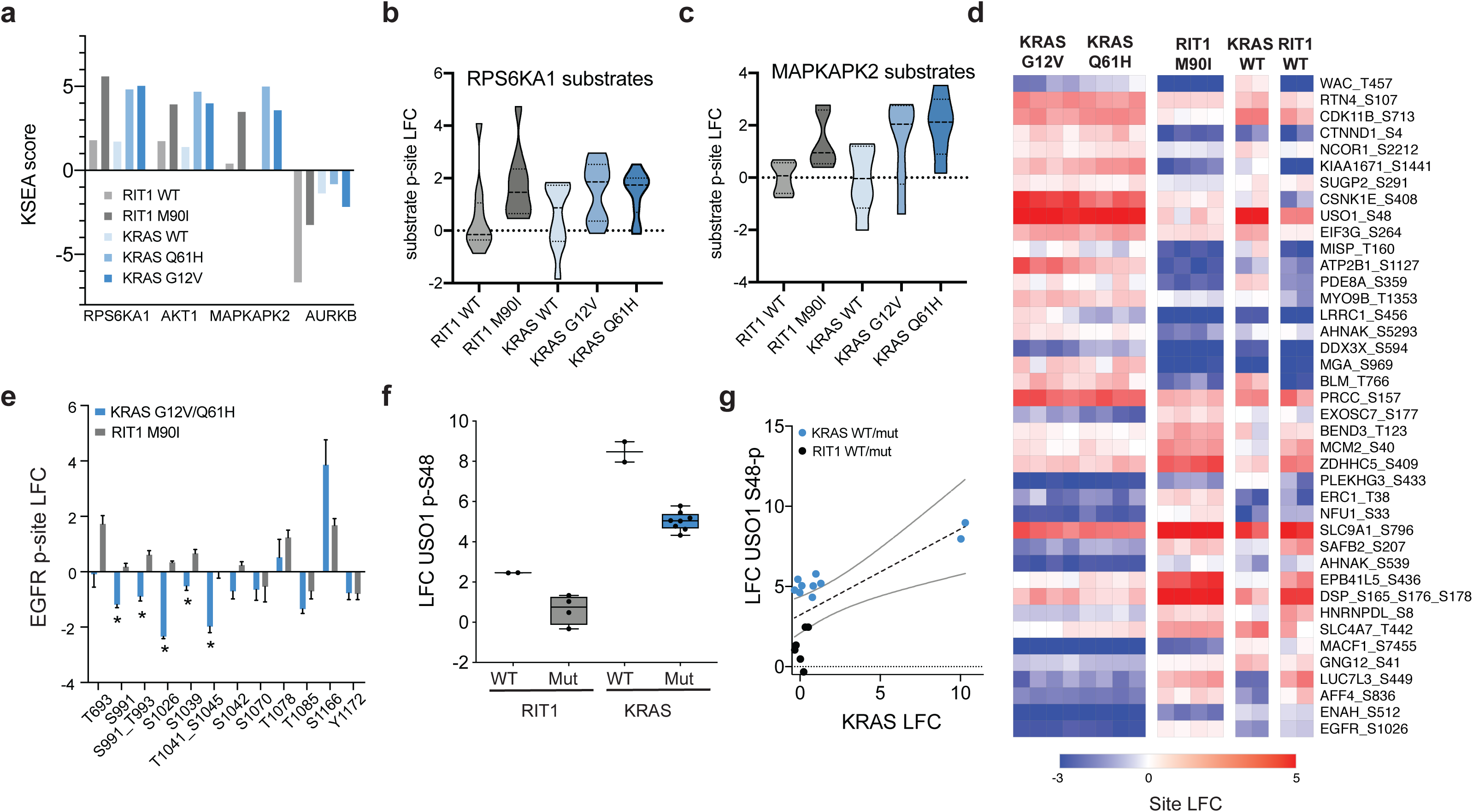
Phosphoproteomic profiling illuminates novel differential post-translational modifications in RIT1^M90I^- and KRAS-mutant cells. **a**, KSEA analysis of AALE phosphoproteomes. Top differentially phosphorylated kinase substrates are shown. The full KSEA results are shown in **Supplementary Fig 4b-c. b**, Violin plot of phospho-site abundance of phospho-sites that are RPS6KA1 substrates. **c**, Violin plot of phospho-site abundance of phospho-sites that are MAPKAPK2 substrates. **d**, Marker selection analysis identifies differentially phosphorylated sites in KRAS-mutant cells compared to RIT1-mutant cells. Phosphosites from KRAS-mutant and RIT1^M90I^-mutant replicate-level phosphoproteome profiles (**Supplementary File 2**) were compared by two-tailed t-test. The top 20 significantly (FDR < 0.05) differentially phosphorylated sites in each direction are shown and ranked by t-statistic. A heat map displays the LFC in phosphorylated peptide abundance of each site compared to vector control, after normalizing to total protein abundance. **e**, LFC of EGFR phosphosites in KRAS-mutant and RIT1-mutant cells. Data shown is the mean + SD of n=8 KRAS-mutant replicates and n=4 RIT1-mutant replicates. *, FDR < 0.01 as determined by two-tailed t-test and two-stage linear step-up procedure of Benjamini, Krieger and Yekutieli. **f**, Relative phosphorylation of USO1 at serine 48 as determined by LC-MS/MS. Box and whiskers show the 25th-75th percentiles and minimum to maximum of the data, respectively. **g**, Relationship of USO1 S48 phosphorylation to KRAS total protein abundance. A dashed line displays the linear regression fit and gray lines display the 95% confidence interval of the linear model. *r* = 0.70, *p* < 0.01.

Next we assessed the divergent functions of RIT1^M90I^ and KRAS^G12V^/KRAS^Q61H^ by identifying proteins with differential phosphorylation in KRAS-mutant versus RIT1^M90I^-mutant cells. 902 differentially phosphorylated sites were identified by two-tailed t-test and multiple hypothesis correction (**Fig. 5d; FDR < 0.05).** Interestingly, the top site with lower phosphorylation in KRAS^G12V^ and KRAS^Q61H^ cells was EGFR serine 1026 (**Fig. 5d**). In lung adenocarcinoma, *KRAS* mutations and *EGFR* mutations are mutually exclusive, suggesting a powerful genetic interaction between these two genes. Recent work demonstrated that mutant KRAS and EGFR display synthetic lethality^37^. However the mechanism underlying this lethality is unknown. Further inspection of the phospho-proteome signatures revealed extensive alteration of EGFR phosphorylation by KRAS^G12V^ and KRAS^Q61H^, but not by RIT1^M90I^. 11 of 12 EGFR sites detected by LC-MS/MS occur in the cytoplasmic carboxy-terminal tail of EGFR (**Fig. 5e**). Five of these sites (S991, S991/T993 double phosphorylation, S1026, S1039, and T1041/S1045 double phosphorylation) were significantly depleted of phosphorylation in KRAS^G12V^ and KRAS^Q61H^-expressing cells but not in RIT1^M90I^-expressing cells. Interestingly, these sites lie in a region of EGFR that is involved in receptor internalization and endocytosis^38^ and a phosphorylation-deficient mutant at S991 is defective at internalization^39^. Consistently, EGFR protein abundance was increased in KRAS-mutant cells (**Supplementary Fig. 4g**) Although the specific regulatory mechanisms leading to this depletion remain unknown, these data point to the existence of feedback regulatory signaling from oncogenic KRAS to EGFR.

Examining phosphorylation uniquely promoted by KRAS^G12V^ and KRAS^Q61H^, we identified USO1 phosphorylation at S48 as one of the top most significantly increased phosphorylation events in KRAS-mutant cells. USO1, also known as p115, is a vesicle tethering factor involved in ER-Golgi intracellular trafficking^40^. Although wild-type KRAS and KRAS-mutant proteomic signatures were largely divergent, USO1 serine 48 phosphorylation was promoted by both KRAS^WT^ and mutant KRAS (**Fig. 5f**). KRAS relies on vesicle trafficking to ensure proper post-translational farnesylation and palmitoylation, which are required for targeting of KRAS to the plasma membrane^41^. We hypothesized that USO1 S48 phosphorylation was therefore correlated with KRAS expression rather than activity. Indeed, a significant correlation was observed between overall KRAS expression and USO1 phosphorylation (**Fig. 5g**). In contrast, USO1 S48 phosphorylation was only modestly changed in RIT1-mutant cells (**Fig. 5d**). Notably, RIT1 lacks the farnesylation and palmitoylation signals present in RAS isoforms^42^, so the differential regulation of USO1 by KRAS and RIT1 may be related to differences in RIT1 and KRAS trafficking.

Also among the top differentially phosphorylated sites were 32 phosphorylation sites in AHNAK proteins 1 and 2. AHNAK and AHNAK2 are large scaffolding proteins that have been implicated as tumor suppressor proteins in breast and lung cancer^43,44^. Among all phospho-proteins, a higher proportion (32/117) of sites on AHNAK and AHNAK2 were differentially phosphorylated than expected by chance (P < 0.0001 by Chi Square test; **Supplementary Fig. 4h**). Intriguingly, two recent proximity-labeling proteomic studies identified AHNAK and AHNAK2 as KRAS-interacting proteins^45,46^, raising the possibility that a direct physical interaction between KRAS and AHNAK proteins may be involved in the differential AHNAK phosphorylation we observe.

## Discussion

Here we describe quantitative proteomic, phosphoproteomic, and transcriptomic datasets that provide the first systematic view of the RIT1^M90I^-regulated signaling network. These datasets were generated from isogenic human lung epithelial cells to provide a physiological view of the consequences of RIT1 activation in the same cellular compartment that is involved in lung adenocarcinoma, a tumor type with prevalent mutations in *KRAS* and *RIT1*. Broadly, we find that ‘omic signatures from RIT1^M90I^-expressing cells largely phenocopy those from cells with overexpression of wild-type RIT1. This finding lends further support to the notion that oncogenic RIT1 variants function at least in part through increasing RIT1 abundance^11^. This is in contrast to KRAS, where overexpression of wild-type KRAS induces signatures unrelated or opposite to that of oncogenic KRAS variants G12V and Q61H. The opposing functions of wild-type and mutant KRAS is consistent with recent evidence suggesting that KRAS functions as a dimer and that wild-type KRAS directly inhibits the function of oncogenic KRAS variants via physical dimerization^47^. This divergence in the function of wild-type RIT1 and KRAS hints at fundamental differences in molecular regulation of each wild-type GTPase. The ability of RIT1 to promote downstream RAF/MEK/ERK signaling when aberrantly expressed suggests that RIT1 may not be subject to the same tight regulation by GTPase-activating proteins (GAPs) that normally keep RAS in an inactive state. Furthermore, these data raise the possibility that wild-type RIT1 overexpression in *RIT1*-amplified cancers may contribute to tumorigenesis. *RIT1*, on chromosome 1q, is frequently amplified in uterine carcinosarcoma, liver hepatocellular cancer, cholangiocarcinoma, breast cancer, lung adenocarcinoma, and ovarian cancer. RIT1 mRNA expression is increased in amplified cases, regardless of tissue type, raising the possibility that RIT1 overexpression could play a role in tumorigenesis in these cancers.

We find that RIT1^M90I^, KRAS^G12V^, and KRAS^Q61H^ share the ability to activate canonical RAS effector pathways PI3K/AKT and RAF/MEK. Likely as a consequence of RAF/MEK signaling to FOSL1, RIT1^M90I^ also shares the ability to induce EMT markers including Vimentin, N-cadherin, and fibronectin. KRAS and RIT1 variants also shared the ability to profoundly suppress HLA-A, - B, and -C expression via a posttranscriptional mechanism. Taking advantage of differential expression of HLA proteins between RIT1^WT^ and all other isogenic lines, we identified an entire Rab-mediated endocytic network that was lost together with HLA proteins in RIT1- and KRAS-mutant cells. This downregulated module also included PSMB9, a subunit of the immunoproteasome that is involved in antigen processing for class I MHC presentation. RAS oncogenes have long been recognized to suppress surface MHC expression^48^, in some cases transcriptionally and in others post-transcriptionally^31^. Our data link both oncogenic RIT1 and RAS to modulation of the processing and trafficking of MHC Class I molecules. Further identification of the mechanism of RIT1/RAS-mediated MHC suppression will provide a better understanding of tumor immune evasion which is critically needed to optimize patient stratification of cancer immunotherapy.

In addition to the largely concordant regulation of proteins by mutant RIT1 and KRAS, we identified several unique phosphoproteins with differential abundance in RIT1^M90I^ and KRAS-mutant cells. These included EGFR, a key oncoprotein in lung adenocarcinoma, which showed reduced phosphorylation of sites involved in receptor internalization and endocytic trafficking. Given the potent genetic interactions between KRAS and EGFR in lung cancer and colon cancer, it is attractive to speculate that feedback regulation of KRAS to EGFR could provide an explanatory mechanism for this phenomenon. Future work is needed to determine the basis of the specific regulation of EGFR phosphorylation by oncogenic KRAS but not RIT1.

Together, these results demonstrate the power of quantitative proteomics and transcriptomics to provide global views of cancer oncogene signaling. Our multi-omic analysis validated known consequences of RAS activation such as EMT and activation of RAF/MEK and PI3K signaling. For the first time, we gained a global view of RIT1 function, which confirmed its ability to stimulate canonical RAS signaling. However, phosphoproteomic profiling identified a number of key divergent mechanisms between KRAS- and RIT1-mutant cells, which point to the existence of novel, unique regulators or effectors of KRAS and RIT1 still to be identified. Future work is needed to investigate the mechanisms of these differences between KRAS and RIT1, the results of which will have important implications for cancer therapy and Noonan Syndrome.

## Methods

### Isogenic Cell Line Generation

Plasmid constructs were cloned using Gateway Technology (Invitrogen/ThermoFisher) using pLX301 destination vector (Broad Institute) and pDONR223-RIT1 donor vectors previously described^5^. Lentivirus was generated by transfection of HEK293T cells with packaging and envelope vectors using standard protocols. AALE cells were a kind gift of Jesse Boehm (Broad Institute). Isogenic cells were generated by transduction of lentivirus generated from pLX317-Renilla luciferase or pLX301-RIT1^WT^, pLX301-RIT1^M90I^, pLX301-KRAS^WT^, pLX301-KRAS^G12V^, or pLX301-KRAS^Q61H^ and selection with puromycin. Stable pools of cells were maintained in small airway growth medium (Lonza).

### Soft Agar Assay

1×10^5^ cells were suspended in 1 ml of 0.33% select agar in small airway growth medium without EGF (Lonza) and plated on a bottom layer of 0.5% select agar in the same media in six-well dishes. Each cell line was analyzed in triplicate. Colonies were photographed after 14–21 days and quantified using CellProfiler^49^.

### Transcriptome profiling

Three technical replicates per cell line were harvested at ∼90% confluence (n = 18 total dishes). Cells were lysed and total RNA was extracted using Direct-zol RNA Miniprep plus (Zymo Research). Libraries were constructed using the non-strand-specific poly-A selection Illumina TruSeq kit for 50bp paired-end reads. Libraries were then pooled and sequenced on the Illumina NovaSeq platform (Fred Hutch Genomics Core). Reads were aligned to the human reference genome build hg19/GRCh37 using STAR v.2.5.3a^50^. Alignments were annotated for duplicates and read groups, and then reordered and indexed, using Picard Tools v.1.114^51^. Read statistics for each RNA-seq sample were calculated using RSeQC^52^. Quantification of gene transcripts was performed by the featureCounts program within the Subread package^53^, using hg19 gene annotation from UCSC Gene level CPM and RPKM values were calculated with edgeR v.3.22.3^54^, and converted into transcripts per million (TPM values with an in-house script. In total, 12,887 genes were identified with average logCPM at least 0.1 across all samples. Differential expression analyses comparing KRAS or RIT1 perturbed cell lines against vector control lines were performed using edgeR^54^.

### High performance liquid chromatography tandem mass spectrometry (LC-MS/MS)

Cells were washed in ice-cold PBS, scraped into PBS, pelleted, and snap frozen in liquid nitrogen. The experimental workflow for sample processing, TMT-labeling, peptide enrichment, and LC-MS/MS were largely as previously described^55^. Briefly, pellets were lysed in 200 µl of chilled urea lysis buffer (8 M urea, 75 mM NaCl, 50 mM Tris (pH 8.0), 1 mM EDTA, 2 µg/ml aprotinin, 10 µg/ml leupeptin, 1 mM PMSF, 1:100 (vol/vol) Phosphatase Inhibitor Cocktail 2, 1:100 Phosphatase Inhibitor Cocktail 3, 10 mM NaF, and 20 µM PUGNAc) for each ∼50 mg portion of wet-weight tissue. Lysates were reduced with 5mM DTT, alkylated with 10 mM IAM, and digestion performed in solution with 1 mAU LysC per 50 µg of total protein and trypsin at an enzyme/substrate ratio of 1:49. Reactions were quenched with FA and brought to pH = 3 with FA. Peptides were desalted on 200 mg tC18 SepPak cartridges and dried by vacuum centrifugation. 340 µg of peptides were labeled with 10-plex Tandem Mass Tag reagents (TMT10, Fisher Scientific), according to manufacturer’s instructions. To enable quantification of peptides across all 12 samples, the samples were labeled in sets of 10 across two different TMT10 pools in a crossover design with 8 of 12 samples analyzed in both TMT10 pools. A 50/50 mix of both AALE vector control lysates was used as an internal reference in both TMT10 runs (**Supplementary Fig. 1b**).

Each TMT10-plex was desalted in a 200 mg tC18 SepPak cartridge and fractionated using offline HPLC. 5% of each fraction was collected into an HPLC vial for proteome analysis by LC-MS/MS. The remaining 95% was processed for phospho-peptide enrichment via immobilized metal affinity chromatography (IMAC). IMAC enrichment was performed using Ni-NTA Superflow Agarose beads incubated with peptides solubilized in a final concentration of 80% MeCN/0.1% TFA. Phospho-enriched peptides were desalted and collected into an HPLC vial for analysis by LC-MS/MS.

Online fractionation was performed using a nanoflow Proxeon EASY-nLC 1200 UHPLC system (Thermo Fisher Scientific) and separated peptides were analyzed on a benchtop Orbitrap Q Exactive Plus mass spectrometer (Thermo Fisher Scientific) equipped with a nanoflow ionization source (James A. Hill Instrument Services, Arlington, MA). In-house packed columns (20 cm x 75 μm diameter C18 silica picofrit capillary column; 1.9 μm ReprosIl-Pur C18-AQ beads, Dr. Maisch GmbH, r119.aq; Picofrit 10 μm tip opening, New Objective, PF360-75-10-N-5). Mobile phase flow rate was 200 nL/min, comprised of 3 % acetonitrile/0.1 % formic acid (Solvent A) and 90 % acetonitrile /0.1 % formic acid (Solvent B). The 110 min LC-MS/MS method consisted of a 10 min column-equilibration procedure; a 20 min sample-loading procedure; and the following gradient profile: (min: % B) 0:2; 2:6; 85:30; 94:60; 95:90; 100:90; 101:50; 110:50 (the last two steps at 500 nL/min flow rate). Data-dependent acquisition was performed using Xcalibur QExactive v2.4 software in positive ion mode at a spray voltage of 2.00 kV. MS1 Spectra were measured with a resolution of 70,000, an AGC target of 3e6 and a mass range from 300 to 1800 m/z. Up to 12 MS/MS spectra per duty cycle were triggered at a resolution of 35,000, an AGC target of 5e4, an isolation window of 0.7 m/z, a maximum ion time of 120 msec, and normalized collision energy of 30. Peptides that triggered MS/MS scans were dynamically excluded from further MS/MS scans for 20 sec. Charge state screening was enabled to reject precursor charge states that were unassigned, 1, or >6. Peptide match was set to preferred for monoisotopic precursor mass assignment.

### Protein-peptide identification, phosphosite localization, and quantification

MS data was interpreted using the Spectrum Mill software package v6.0 pre-release (Agilent Technologies, Santa Clara, CA. MS/MS spectra were merged if they were acquired within +/- 45 sec of each other with the same precursor m/z. Also, MS/MS spectra that did not have a sequence tag length > 0 (i.e., minimum of two masses separated by the in chain mass of an amino acid) or did not have a precursor MH+ in the range of 750-6000 were excluded from searching. MS/MS spectra searches were performed against a concatenated UniProt human reference proteome sequence database containing 58,929 human proteins including isoforms (obtained 10/17/2014) and 150 additional common laboratory contaminants. ESI-QEXACTIVE-HCD-3 scoring parameters were used for both whole proteome and phosphoproteome datasets. Spectra were allowed +/- 20 ppm mass tolerance for precursor as well as product ions, 30% minimum matched peak intensity, and “trypsin allow P” was set as enzyme specificity with up to 4 missed cleavages allowed. Carbamidomethylation at cysteine was set as fixed modification together with TMT10 isobaric labels at lysine residues (N-termini would be considered regardless if it was TMT labelled). Acetylation of protein N-termini andoxidized methionine were set as variable modifications with a precursor MH+ shift range of −18 to 64 Da for the proteome searches. For the phosphoproteome searches the precursor MH+ shift range was set to 0 to 272 Da and variable modifications of phosphorylation of serine, threonine, and tyrosine. Identities interpreted for individual spectra were automatically designated as confidently assigned using the Spectrum Mill autovalidation module to use target-decoy based false discovery rate (FDR) estimates to apply score threshold criteria. For the whole proteome datasets, thresholding was done at the spectral (< 1.2%) and protein levels (< 0.1%). For the phosphoproteome datasets, thresholding was done at the spectral (< 1.2%) and phosphosite levels (< 1.0%).

Replicates across TMT-plexes were highly correlated (**Supplementary Fig. 1c)** with median Pearson *r* = 0.87 and 0.69 for proteome and phosphoproteome, respectively. Technical replicates and biological replicates were merged to generate final total proteome and phospho-proteome profiles for each isogenic cell line (**Supplementary Tables 1 and 2**). Replicate-level profiles are also supplied as JavaScript Object Notation (.json) files that can be visualized and analyzed using the Morpheus Matrix Visualization and Analysis Software at https://software.broadinstitute.org/morpheus (**Supplementary Files 1 and 2**). Differential protein and phospho-site signatures were generated by computing the mean log_2_(fold change) of the abundance of each site in each sample compared to the vector control cells. Statistical significance of differentially abundant proteins and phosphosites was determined by performing a one sample moderated t-test with multiple hypothesis correction (**Supplementary Tables 1 and 2**).

### Integrative Analysis

Correlation of changes in protein expression and changes in RNA expression was modeled using R’s lm() function. 95% prediction intervals were calculated to determine genes with weak concordance between protein and RNA expression.

### Gene Set Enrichment Analysis

Analysis of enrichment of KRAS signaling in differential RNA expression profiles was performed in R with the goseq package^56^. KRAS signaling gene sets were taken from MSigDB hallmark gene sets^22,57^.

### Transcription Factor Target Enrichment Analysis

Analysis of over-representation of Transcription Factor targets was performed with ChEA3 by submitting lists of differentially expressed genes (|LFC| > 1 and FDR < 0.05). ChEA3 performs Fisher’s Exact Test to compare the input gene set to TF target gene sets in six different libraries^58^. Analysis of the Enrichr Queries library was selected as the focus of the present study. Transcription factors resulting from this analysis were annotated as one of four groups of EMT association. These four groups were the Snail gene family, confirmed EMT genes defined by dbEMT^59^, genes shown to be associated with EMT in at least one study in literature, and genes unrelated to EMT.

### Antibodies and immunoblotting

Antibodies against FOSL1 (D80B4), TXNIP (D5F3E), and Vimentin (D21H3) were purchased from Cell Signaling Technology. Vinculin (V9264) was purchased from Sigma Aldrich. Secondary antibodies StarBright Blue 700 Goat anti-Rabbit IgG, StarBright Blue 520 Goat anti-Rabbit IgG and StarBright Blue 520 Goat anti-Mouse IgG (12005867) were purchased from Bio-Rad. Antibody against RIT1 (#53720) was purchased from Abcam. Cell lysates were prepared in RTK lysis buffer with protease (11836153001, Roche) and phosphatase (04906837001, Roche) inhibitors added and quantified by the BCA assay (Thermo Scientific Waltham, MA). Samples were then boiled in Laemmli buffer (1610747, Bio-Rad, Hercules, CA) and 50 ug of protein was loaded onto 4-15% Mini-Protean TGX (4561084, Bio-Rad, Hercules, CA) gels. Protein gels were run and transferred to PVDF membranes (1704274, Bio-Rad, Hercules, CA) according to manufacturer’s instructions. Proteins were detected by specific primary antibody and secondary antibody then visualized using the ChemiDoc MP Imaging System (Bio-Rad, Hercules, CA).

### KSEA analysis

Kinase-substrate enrichment analysis (KSEA)^60^ was performed using the KSEA App^34^ (https://casecpb.shinyapps.io/ksea/) using kinase-substrate mappings from PhosphoSitePlus^35^ and a p value threshold of < 0.05. A minimum of five detected phospho-site substrates were needed for kinases to be included in the analysis. The full list of kinase scores and number of substrates are shown in Supplementary Table 5. 36 kinases had sufficient substrate sites detected to be included in the analysis. Kinase-substrate mappings are shown in Supplementary Table 5.

## DATA AVAILABILITY

The RNA-seq data have been deposited in the NCBI Gene Expression Omnibus database with accession number GSE146479. All mass spectra contributing to this study can be downloaded in the original instrument vendor format from the MassIVE online repository (Accession number to be updated prior to publication.)

## ACKNOWLEDGEMENTS

We thank Drs. Athea Vichas and Jon Cooper (Fred Hutchinson Cancer Research Center) for advice, discussion, and critical reading of the manuscript. We thank Dr. D.R. Mani (Broad Institute) for guidance on statistical analysis. This research was funded in part through the National Cancer Institute (NCI) K99/R00 CA197762 to AHB, NIH/NCI Cancer Center Support Grant P30 CA015704, NCI Clinical Proteomic Tumor Analysis Consortium grants NIH/NCI U24-CA210986 and NIH/NCI U01 CA214125 to SAC. AL was supported in part by NSF IGERT DGE-1258485. KH was supported in part by PHS NRSA T32GM007270 from NIGMS.

## AUTHOR CONTRIBUTIONS

A.H.B. conceived of and directed the study. S.C. and P.M. supervised the LC-MS/MS experiments. F.M. performed the proteomics data analysis, with contributions from A.H.B. and K.H. A.L. performed the transcriptome analysis. A.L. and A.H.B. wrote the manuscript. A.L., K.H., S.M., I.F., S.F., J.W., and A.H.B. performed experiments. All authors discussed results and provided input on the manuscript.

## SUPPLEMENTARY FIGURE LEGENDS

**Supplementary Figure 1. Workflow and quality control of proteomic and transcriptomic profiling. a**, Replicate-level workflow for tandem mass tag (TMT) labeling and LC-MS/MS. Lysates from duplicate sets of six isogenic cell lines were used to generate two TMT-plex sets, with control samples used to link the two sets. **b**, TMT 10-plex layout showing mass tags associated with each replicate. **c**, Average pairwise replicate correlations (Pearson *r*) of all replicates from each sample group indicated. **d**, Enrichment analysis of differentially expressed genes between KRAS or RIT1 perturbed lines and vector controls using *goseq*^*56*^. mSigDB hallmark gene sets specific to KRAS signaling are shown. **e**, RNA-seq run and mapping statistics show total reads, mapped reads, and reads mapped to rRNA, for each sample.

**Supplementary Figure 2. RIT1 and KRAS promote epithelial-to-mesenchymal transition. a**, Changes in mRNA transcript levels of EMT genes *VIM, CDH2, FN1*, and *KRT19*, in each isogenic cell line compared to vector control. LFC, log_2_(fold-change) compared to vector cells. **b**, Transcription factor target enrichment analysis using Enrichr libraries of differentially expressed genes in RIT1^WT^, KRAS^WT^, and KRAS^Q61H^-mutant cells. FET, Fisher’s exact test. Red = Snail family. Orange = confirmed EMT genes in dbEMT^59^. Pale orange = EMT-associated genes in literature.

**Supplementary Figure 3. Post-transcriptional loss of Class I MHC proteins. a**, Rank plot of all protein abundance changes in KRAS^Q61H^-mutant or RIT1^M90I^-mutant cells compared to vector control, generated by LC-MS/MS. HLA-A,-B,-C, and -F proteins are labeled. LFC, log_2_ fold-change. **b**, Change in mRNA transcript levels of HLA genes and upstream regulators of MHC Class I, in each isogenic cell line compared to vector controls. LFC, log_2_ fold change compared to vector cells. **c**, Correlation of protein levels in HLA-A and PSMB9 across each isogenic cell line. A line is the best-fit linear regression with significant non-zero slope (p < 0.05).

**Supplementary Figure 4. Phosphoproteome profiling identifies enhanced phosphorylation of specific kinase substrates in KRAS- and RIT1-mutant cells. a**, Pairwise replicate correlation (Pearson *r*) heatmap and unsupervised clustering of phosphoproteome data. **b**, KSEA of phosphoproteome data for RIT1^WT^ and RIT1^M90I^-expressing cells. The kinase z-score indicates the overall score for each kinase listed, normalized by the total number of substrates. Significant scores (p<0.05) are indicated in red and blue. Phospho-sites of kinases in red were more highly abundant in the cell line compared to vector control, whereas phospho-sites of kinases in blue were more highly abundant in vector control than the indicated cell line. **c**, KSEA of phosphoproteome data for KRAS-expressing cells. Labeling as in (**b**). **d**, Violin plot of phospho-site abundance of AKT1 substrate sites. **e**, Violin plot of phospho-site abundance of PRKCA substrate sites. **f**, Violin plot of phospho-site abundance of AURKB substrate sites. **g**, EGFR protein abundance in LC-MS/MS data compared to vector control. **f**, Proportion of phosphorylated sites in AHNAK proteins with differential phosphorylation between KRAS-mutant and RIT1^M90I^-mutant cells. Data shown is the percentage of differentially abundant phosphorylation sites in AHNAK and AHNAK2 compared to all other sites. Significance was determined from the analysis in (**b**), FDR < 0.05. ****, p < 0.0001 by two-sided Fisher’s exact test.

## Notes

### Competing Interest Statement

The authors have declared no competing interest.

