## Supplemental Figures for "Mapping the unique and shared functions of oncogenic KRAS and RIT1 with proteome and transcriptome profiling"

**a**

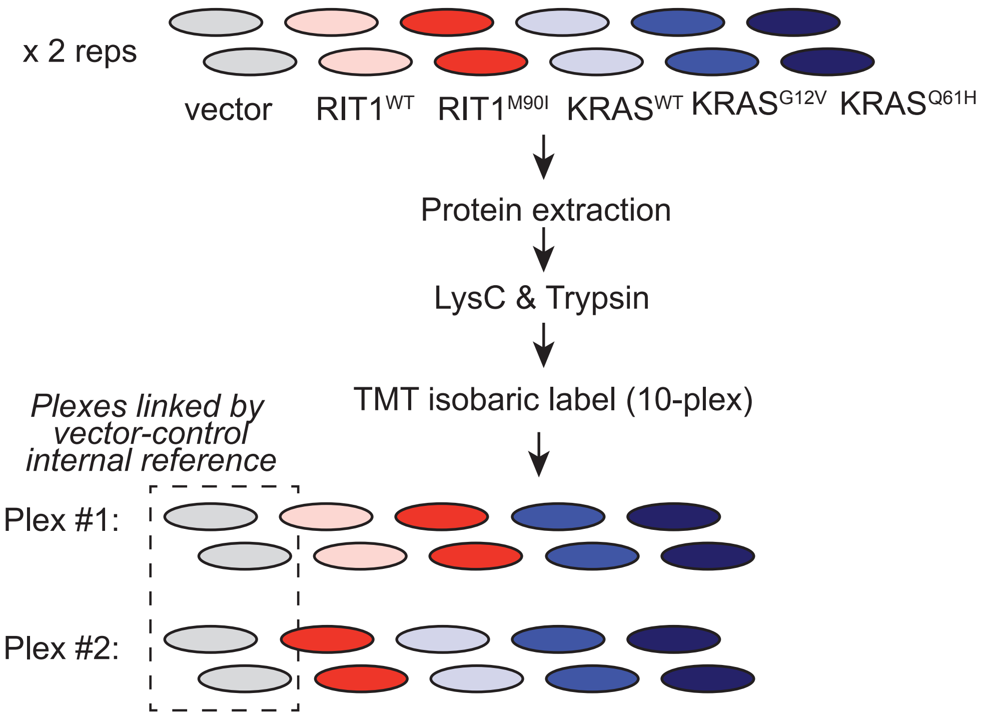


**b**

| *Experiment 1* | *TMT Tag* |
|---|---|
| KRAS G12V Rep 1 | 126 |
| RIT1 WT Rep 2 | 131 |
| RIT1 M90I Rep 2 | 127C |
| KRAS Q61H Rep 2 | 127N |
| Vector Rep 1 | 128C |
| KRAS G12V Rep 2 | 128N |
| RIT1 M90I Rep 1 | 129C |
| RIT1 WT Rep 1 | 129N |
| KRAS Q61H Rep 1 | 130C |
| Vector Rep 2 | 130N |

| *Experiment 2* | *TMT Tag* |
|---|---|
| KRAS G12V Rep 1 | 126 |
| RIT1 M90I Rep 1 | 131 |
| KRAS Q61H Rep 2 | 127C |
| KRAS Q61H Rep 1 | 127N |
| KRAS WT Rep 2 | 128C |
| KRAS WT Rep 1 | 128N |
| Vector Rep 1 | 129C |
| RIT1 M90I Rep 2 | 129N |
| KRAS G12V Rep 2 | 130C |
| Vector Rep 2 | 130N |

**c**

| *Replicate set* | *Proteome* | *Phospho-proteome* |
|---|---|---|
| RIT1 WT | 0.84 | 0.60 |
| RIT1 M90I | 0.83 | 0.69 |
| KRAS WT | 0.90 | 0.77 |
| KRAS G12V | 0.87 | 0.69 |
| KRAS Q61H | 0.91 | 0.72 |

**d**

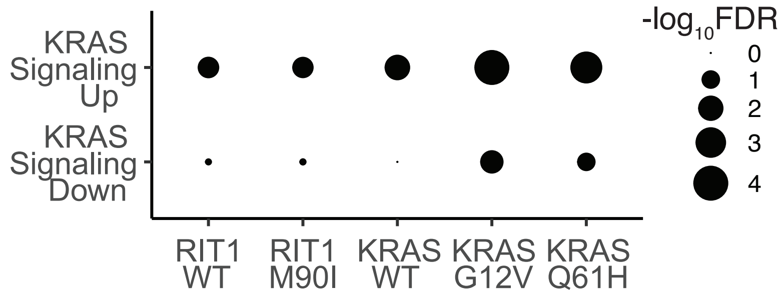


**e**

| Group | Sample | Total Reads | Mapped Reads | rRNA Reads |
|---|---|---|---|---|
| KRAS-G12V | KRAS-G12V_1 | 265969454 | 102506440 | 1413908 |
| KRAS-G12V | KRAS-G12V_2 | 236719962 | 63365494 | 1239580 |
| KRAS-G12V | KRAS-G12V_3 | 246633574 | 77975208 | 1684446 |
| KRAS-Q61H | KRAS-Q61H_1 | 230047062 | 75117724 | 2306602 |
| KRAS-Q61H | KRAS-Q61H_2 | 212706418 | 67632558 | 1454868 |
| KRAS-Q61H | KRAS-Q61H_3 | 212464086 | 80122798 | 719300 |
| KRAS-WT | KRAS-WT_1 | 215832806 | 75204614 | 1927496 |
| KRAS-WT | KRAS-WT_2 | 114503272 | 54598020 | 915628 |
| KRAS-WT | KRAS-WT_3 | 275145008 | 86902056 | 1710894 |
| Vector | Vector_1 | 208952294 | 77102730 | 1373232 |
| Vector | Vector_2 | 250094672 | 79498920 | 2074506 |
| Vector | Vector_3 | 188129714 | 55042564 | 1131944 |
| RIT1-M90I | RIT1-M90I_1 | 219121344 | 53220806 | 2628260 |
| RIT1-M90I | RIT1-M90I_2 | 228132502 | 67300612 | 2385516 |
| RIT1-M90I | RIT1-M90I_3 | 180861518 | 34480460 | 1295320 |
| RIT1-WT | RIT1-WT_1 | 223601164 | 59893560 | 1913980 |
| RIT1-WT | RIT1-WT_2 | 254163922 | 72581670 | 1783556 |
| RIT1-WT | RIT1-WT_3 | 211397706 | 63770336 | 1824880 |

### Supplementary Figure 2

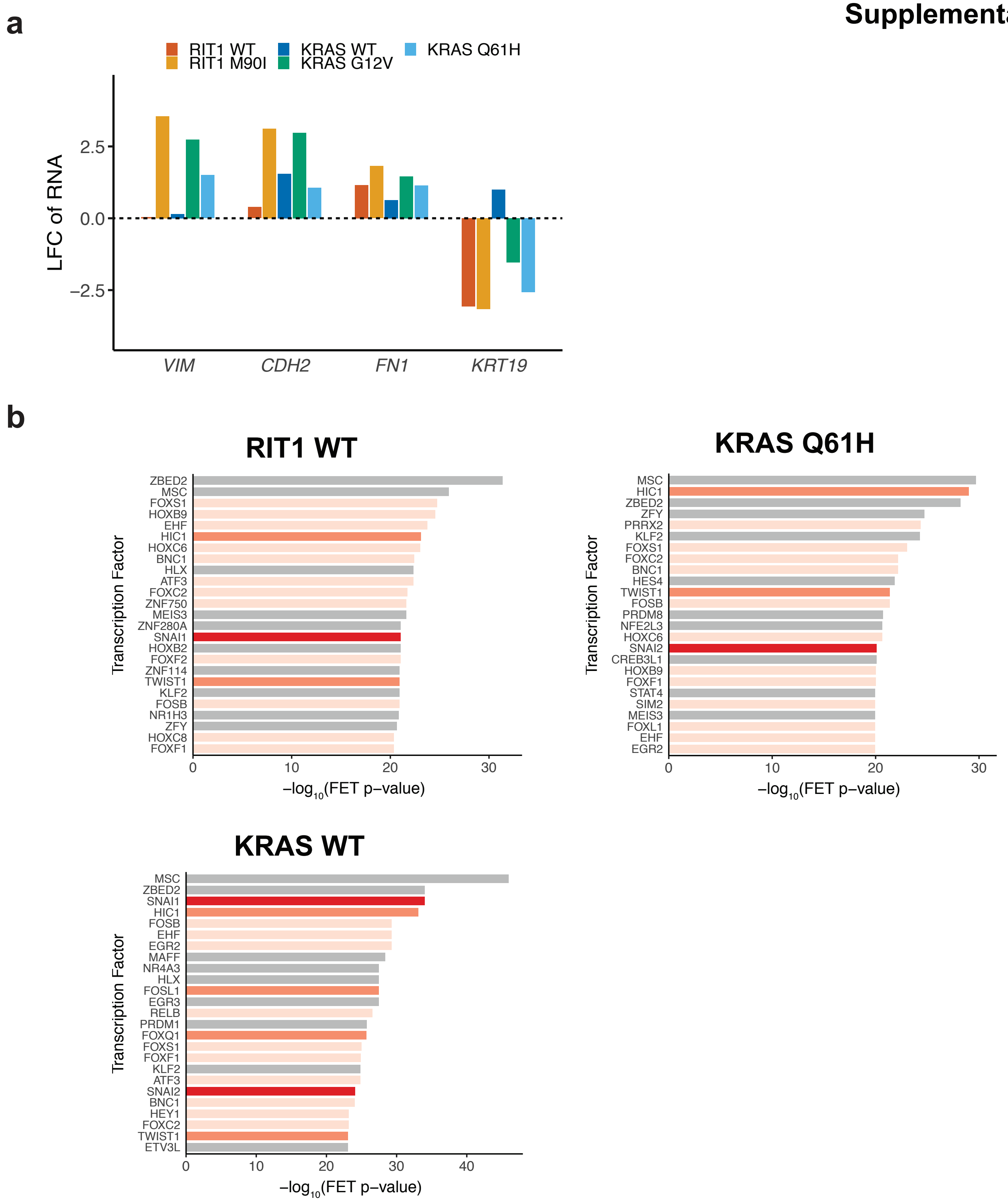

### Supplementary Figure 3

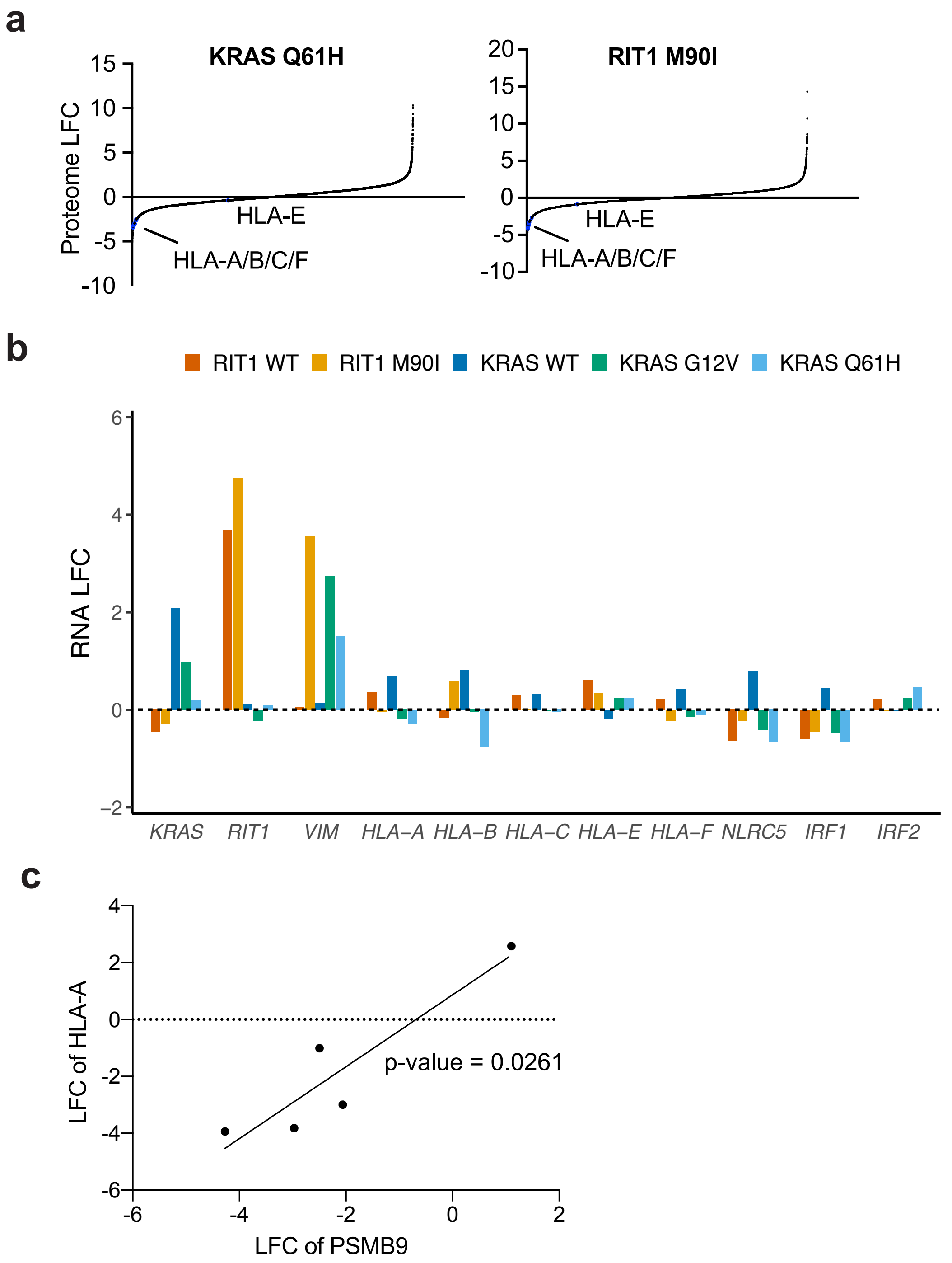

### Supplementary Figure 4

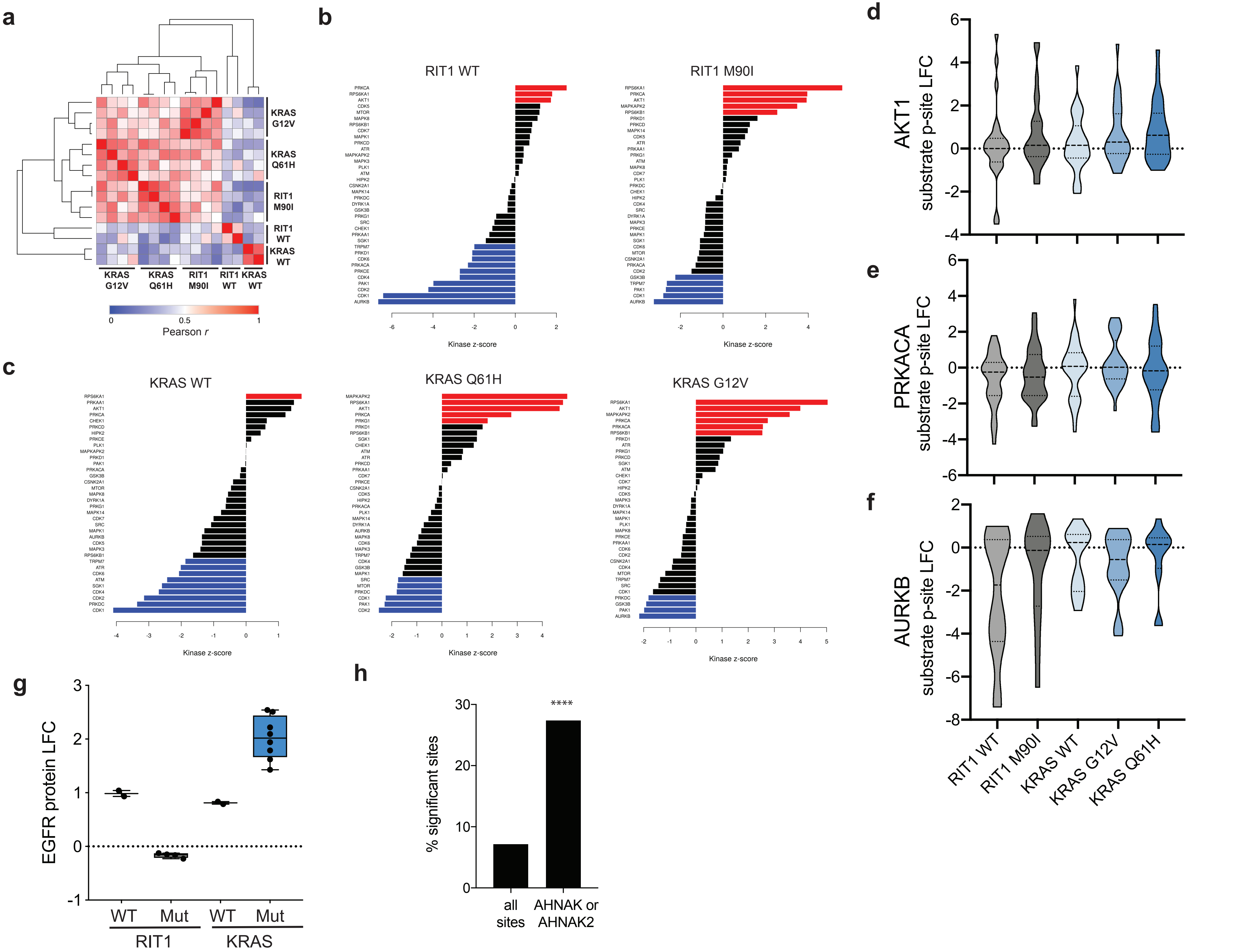
